# Identification of *Candida glabrata* Transcriptional Regulators that Govern Stress Resistance and Virulence

**DOI:** 10.1101/2020.03.10.986703

**Authors:** Elan E. Filler, Yaoping Liu, Norma V. Solis, Luis F. Diaz, John E. Edwards, Scott G. Filler, Michael R. Yeaman

## Abstract

The mechanisms by which *Candida glabrata* resists host defense peptides and caspofungin are incompletely understood. To identify transcriptional regulators that enable *C. glabrata* to withstand these classes of stressors, a library of 215 *C. glabrata* transcriptional regulatory deletion mutants was screened for susceptibility to both protamine and caspofungin. We identified 8 mutants that had increased susceptibility to both host defense peptides and caspofungin. Of these mutants, 6 were deleted for genes that were predicted to specify proteins involved in histone modification. These genes were *ADA2, GCN5, SPT8, HOS2, RPD3*, and *SPP1*. The *ada2*Δ and *gcn5*Δ mutants also had increased susceptibility to other stressors such as H_2_O_2_ and SDS. In the *Galleria mellonella* model of disseminated infection, the *ada2*Δ and *gcn5*Δ mutants had attenuated virulence, whereas in neutropenic mice, the virulence of the *ada2*Δ and *rpd3*Δ mutants was decreased. Thus, histone modification plays a central role in enabling *C. glabrata* to withstand host defense peptides and caspofungin, and Ada2 is essential for the maximal virulence of this organism during disseminated infection.

## Introduction

In the United States of America, *Candida glabrata* causes approximately one third of cases of hematogenously disseminated candidiasis (1, 2). The echinocandin antifungal agents are currently the primary treatment for disseminated candidal infections, especially those caused by *C. glabrata* (3). Of significant concern, echinocandin antifungal drugs are being isolated with increasing frequency (4). In some medical centers, up to 25% of *C. glabrata* isolates are either intermediately susceptible or resistant to echinocandins (5) Many strains of *C. glabrata* are resistant to echinocandins because of mutations in *FKS1* or *FKS2* that encode 1,3 β-glucan synthases, the target enzymes of these drugs (6). However, other strains are resistant to echinocandins by other, unknown mechanisms (2, 7).

Host defense peptides play key roles in the host defense against *C. glabrata* infections. These peptides, such as α-defensins, β-defensins, cathelicidins and LL-37, are produced by leukocytes and the epithelial cells that line cutaneous and mucosal barriers (8, 9). For *C. glabrata* to colonize the host and persist within the deep tissues, it must be able to resist inhibition or killing by these host defense peptides. While some strains of *C. glabrata* are readily killed by β-defensins, other strains are relatively resistant (10). The mechanisms by which *C. glabrata* resists host defense peptides are not fully understood.

To gain a deeper understanding of the mechanisms by which *C. glabrata* resists both echinocandins and host defense peptides, we screened a library of 215 *C. glabrata* transcriptional regulator mutants for strains that were susceptible to the echinocandin, caspofungin and the host defense peptide, protamine. Remarkably, of the 8 strains that were found to susceptible to both compounds, 6 contained mutations in genes whose products are predicted to function in histone modification including acetylation, deacetylation, and methylation. Among these mutants, the Δ*ada2* and Δ*gcn5* mutants were also found to have increased sensitivity to hydrogen peroxide and SDS. Both the Δ*ada2* and the Δ*rpd3* mutants had attenuated virulence during disseminated infection in neutropenic mice. These results indicate that in *C. glabrata*, histone modification is required for the organism to resist caspofungin and protamine, and that Ada2 and Rbd3 govern virulence during disseminated infection.

## Results

### Screening the transcription factor deletion mutant library identified eight genes that governed susceptibility to protamine and caspofungin

In total, 215 *C. glabrata* deletion mutants were screened for susceptibility to protamine and caspofungin. This screen identified 17 mutants that had increased susceptibility to protamine alone and 8 that had increased susceptibility to caspofungin alone, relative to the wild-type strain (Supplemental Table 1). Eight mutants had increased susceptibility to both protamine and caspofungin (Table 1; Fig. 1). Of the genes that were mutated in these eight strains, six specified proteins involved in histone modification. These proteins included Ada2, Gcn5, and Spt8, which are components of the SAGA histone acetyltransferase complex (11), Hos2 and Rpd3, which are members of same family of histone deacetylases (12), and Spp1, a component of the COMPASS histone methyltransferase complex (13). To verify that Ada2, Gcn5, Hos2, Rpd3, and Spp1 governed susceptibility to protamine and caspofungin, we complemented each of the *C. glabrata* mutants with an intact allele of the gene that was disrupted. Complementation restored the wild-type phenotype in all strains (Figure 1). These results indicate that histone modification is the key mechanism by which *C. glabrata* resists host defense peptides and caspofungin.

**Table 1.** Results of the mutant screen. List of genes that were required for resistance to both protamine and caspofungin.

| Systematic Name | Gene Name | Putative Function |
| --- | --- | --- |
| CAGL0K06193g | *ADA2* | Transcriptional coactivator; component of the Spt-Ada-Gcn5 acetyltransferase (SAGA) complex |
| CAGL0F08283g | *GCN5* | Histone acetyltransferase; component of the Spt-Ada-Gcn5 acetyltransferase (SAGA) complex |
| CAGL0F01837g | *SPT8* | Component of the Spt-Ada-Gcn5 acetyltransferase (SAGA) complex |
| CAGL0A03322g | *HOS2* | NAD-dependent histone deacetylase; subunit of the Set3 and Rpd3L complex |
| CAGL0B01441g | *RPD3* | Histone deacetylase; regulates transcription and silencing; subunit of the Rpd3L complex |
| CAGL0M02585g | *SPP1* | Transcriptional activator; histone methyltransferase; part of Set1/COMPASS complex |
| CAGL0M06831g | *CRZ1* | Transcription factor; downstream component of the calcineurin signaling pathway |
| CAGL0J10274g | *MKS1* | Pleiotropic negative transcriptional regulator |

**Fig. 1.**
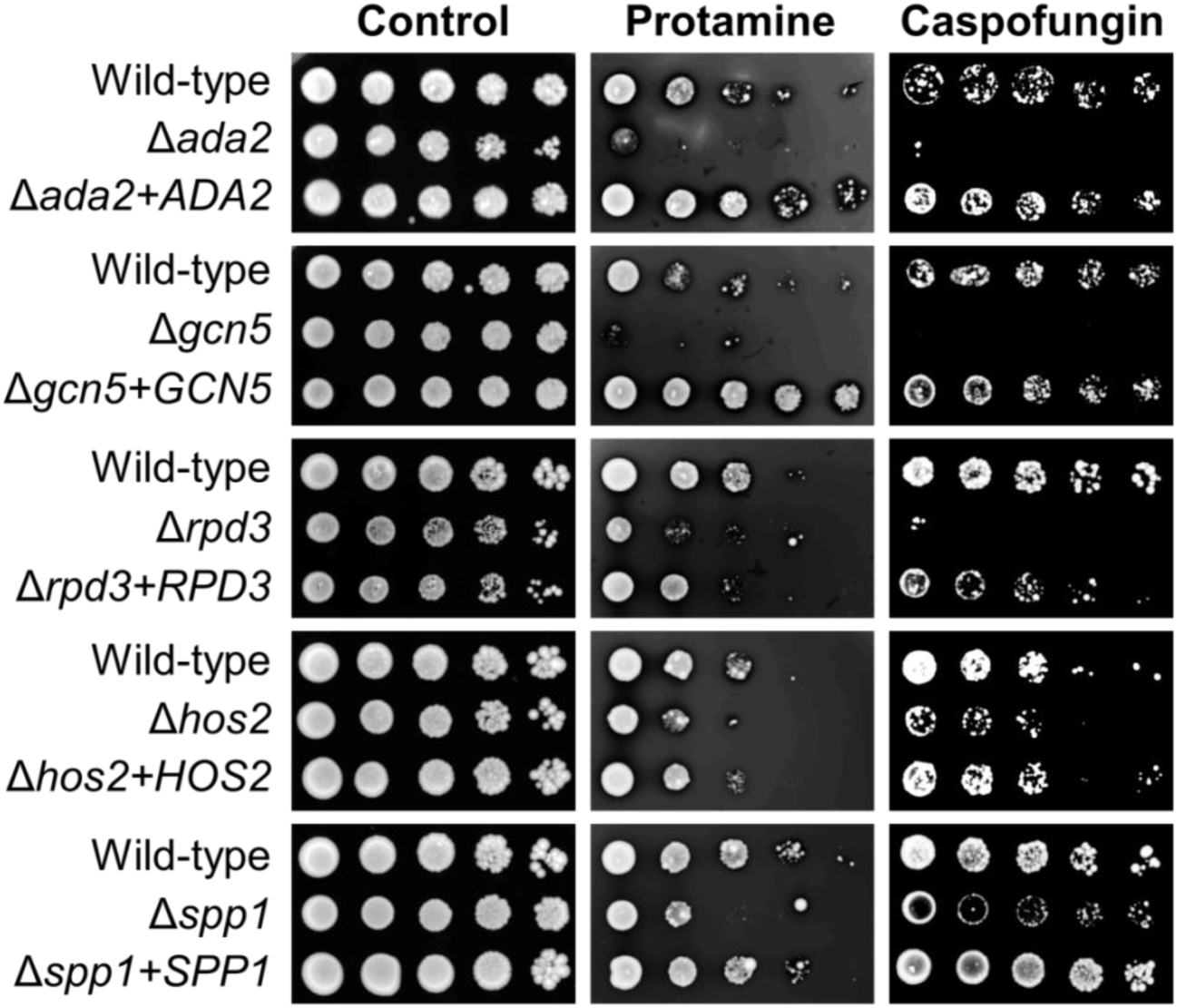
Susceptibility of the various *C. glabrata* mutants to protamine and caspofungin. Serial 10-fold dilutions of the indicated strains were grown on YPD agar alone or on agar containing either protamine or caspofungin at 37°C and then imaged after 48 h.

Two additional mutants, *crz1*Δ and *mks1*Δ, were identified that had increased susceptibility to both protamine and caspofungin. Neither Crz1 nor Mks1 are involved in histone modification (Table 1). Crz1 is a transcription factor in the calcineurin signaling pathway (14), and Mks1 is a negative transcriptional regulator that inhibits multiple signaling pathways such as the target of rapamycin (TOR) and Ras-cyclic AMP pathways (15, 16). These data demonstrate that diverse signaling pathways govern susceptibility to protamine and caspofungin.

### Ada2 and Gcn5 govern resistance to multiple stressors

Next, we evaluated whether histone modification is required for *C. glabrata* to withstand stressors other than protamine and caspofungin. We determined that the *ada2*Δ mutant had increased susceptibility to calcofluor white, H_2_O_2_, SDS and NaCl relative to the wild-type and *ada2*Δ+*ADA2* complemented strains (Fig. 2). The *gcn5*Δ mutant had increased susceptibility to H_2_O_2_ and SDS as compared to the wild-type and *gcn5*Δ+*GCN5* complemented strains. By contrast, the *rpd3*Δ, *hos2*Δ, and *spp1*Δ mutants had wild-type susceptibility to all stressors tested. Thus, the members of the SAGA complex, particularly Ada2, are required for *C. glabrata* to withstand multiple different types of stress.

**Fig. 2.**
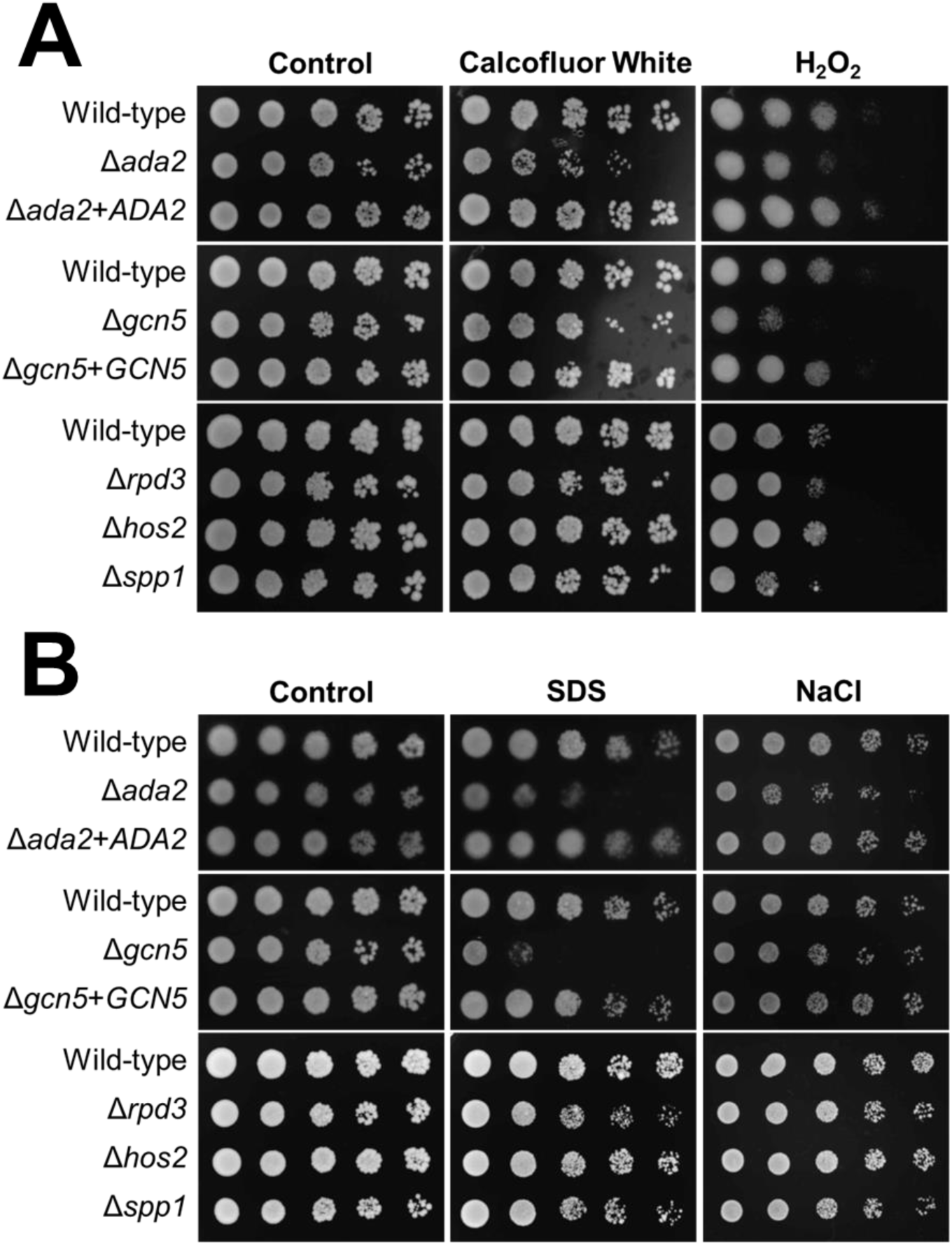
Susceptibility of *C. glabrata* mutants to additional stressors. Serial 10-fold dilutions of the indicated strains were grown on YPD agar alone or on agar containing the indicated stressors at 37°C and then imaged after 48 h.

Protamine was used as a representative host defense peptide in the screening assays. To verify that Ada2 governed the susceptibility of *C. glabrata* to human host defense peptides, we tested the susceptibility of the *ada2*Δ mutant to human neutrophil defensin-1 (hNP-1) and human β-defensin-2 (hBD-2) using a radial diffusion assay. As compared to the wild-type strain and the *ada2*Δ+*ADA2* complemented strains, the *ada2*Δ mutant had increased susceptibility to both of these host defense peptides (Fig. 3).

**Fig. 3.**
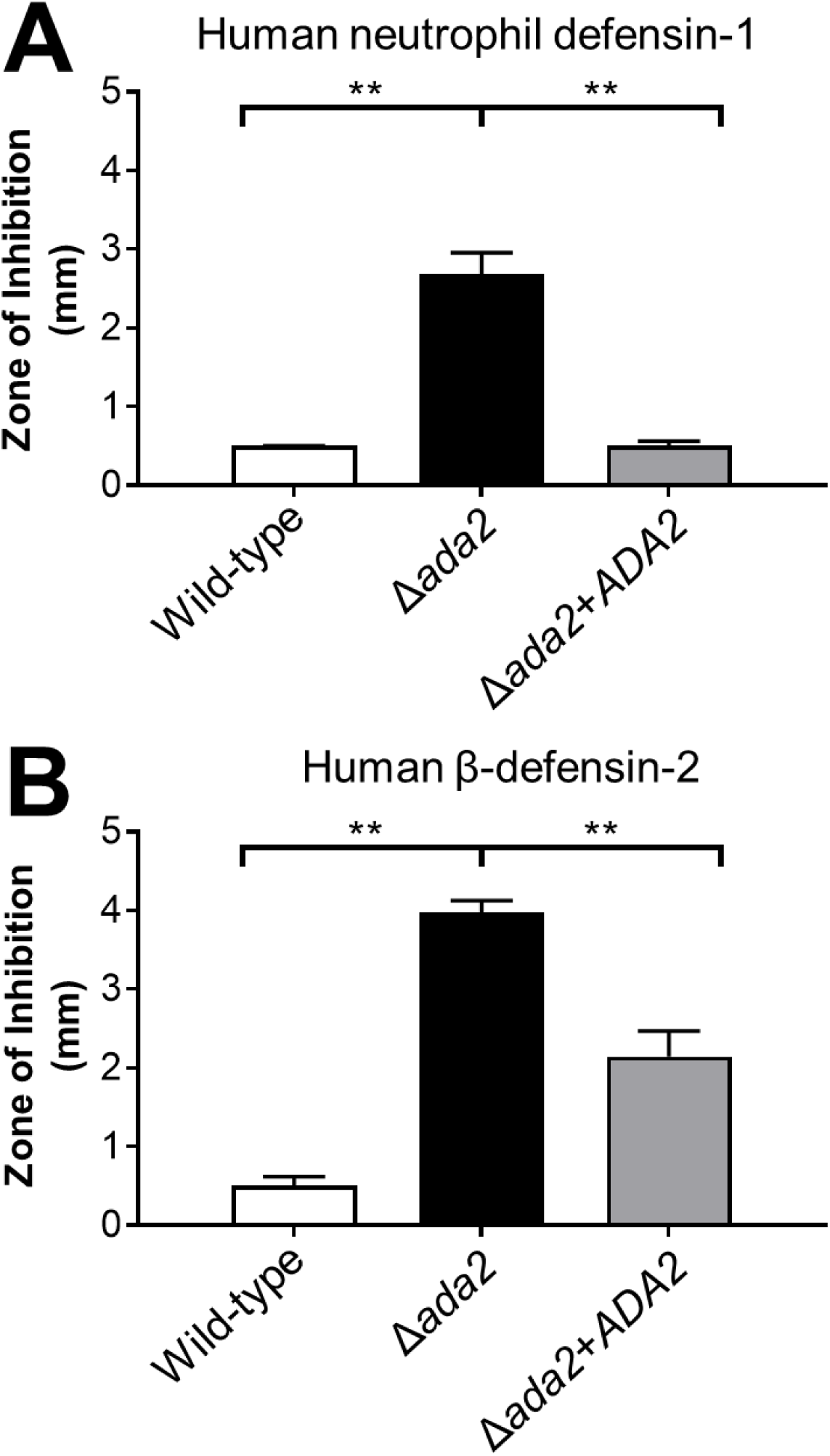
The *ada2*Δ mutant has increased susceptibility to human defensins. The susceptibility of the indicated strains to human neutrophil defensin-1 (hNP-1) (A) and human β-defensin-2 (hBD-2) (B) was determined by radial diffusion assay. Results are the mean ± SD duplicate determinations. \*\**P* < 0.01 by analysis of variance with the Tukey test for multiple comparisons.

### Ada2 is required for virulence

We hypothesized that mutants that were susceptible to both protamine and caspofungin would also have attenuated virulence. To test this hypothesis, we tested the virulence *ada2*Δ, *gcn5*Δ, *rpd3*Δ, *hos2*Δ, and *spp1*Δ mutants in two models of disseminated candidiasis, *G. mellonella* larvae and immunosuppressed mice. In the *G. mellonella* model, each mutant was tested at inocula of 2.5 × 10^6^ and 5 × 10^6^ organisms per larvae. Of the five mutants tested, only the *ada2*Δ and gcn5Δ mutants had virulence defects. Larvae infected with either the high or low inoculum of the Δ*ada2* mutant had significantly longer survival than those infected with the wild-type strain and the *ada2*Δ+ADA2 complemented strain (Fig 4). Larvae infected with the low inoculum of the *gcn5*Δ mutant also survived significantly longer than larvae infected with the controls strain. However, when injected at the high inoculum, the virulence defect of the *gcn5*Δ mutant was no longer apparent. Collectively, these results demonstrate that Ada2, and to a lesser extent Gcn5, are required for the full virulence of *C. glabrata* in *G. mellonella*.

**Fig. 4.**
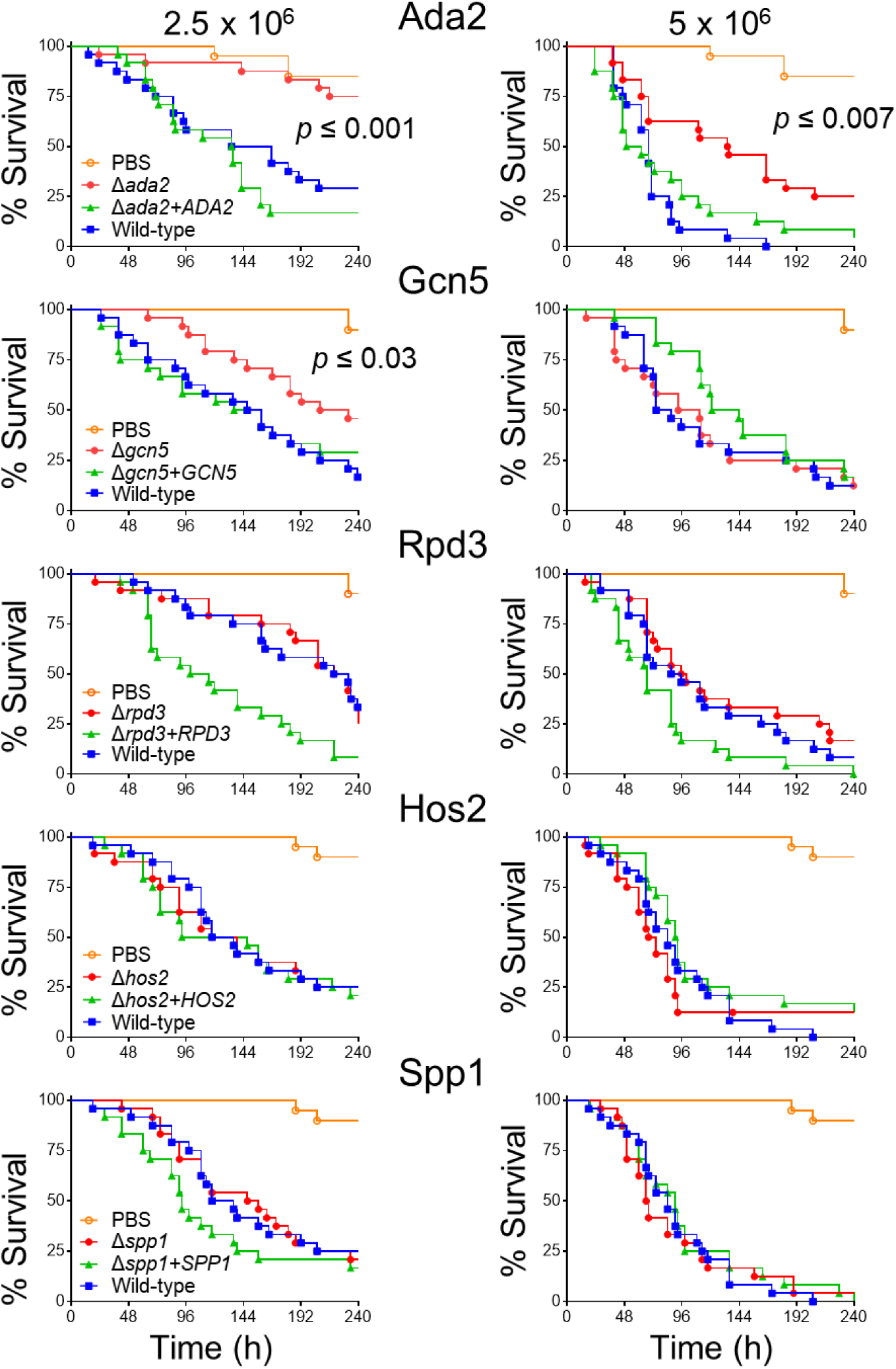
Virulence testing in *G. mellonella*. *G. mellonella* larvae were infected with either 2.5 × 10^6^ cells (left panels) or 5 × 10^6^ (right panel) cells of the indicated strains of *C. glabrata* and then monitored twice daily for survival. Survival curves are the combined results of 2 independent experiments for a total of 24 larvae per strain. The indicated *p* values were determined using the Gehan-Breslow-Wilcoxon test, comparing the larvae infected with the deletion mutant with those infected with the wild-type and complemented strains.

The virulence of the five transcriptional regulatory mutants was also tested in the neutropenic mouse model of disseminated candidiasis using kidney fungal burden as the primary endpoint. Both the *ada2*Δ and *rpd3*Δ mutants had significant virulence defects (Fig. 5). The median kidney fungal burdens of mice infected with these mutants was 50-fold and 80-fold lower than that of mice infected with the wild-type strain. In contrast to its attenuated virulence in *G. mellonella*, the *gcn5*Δ mutant had wild-type virulence in mice.

**Fig. 5.**
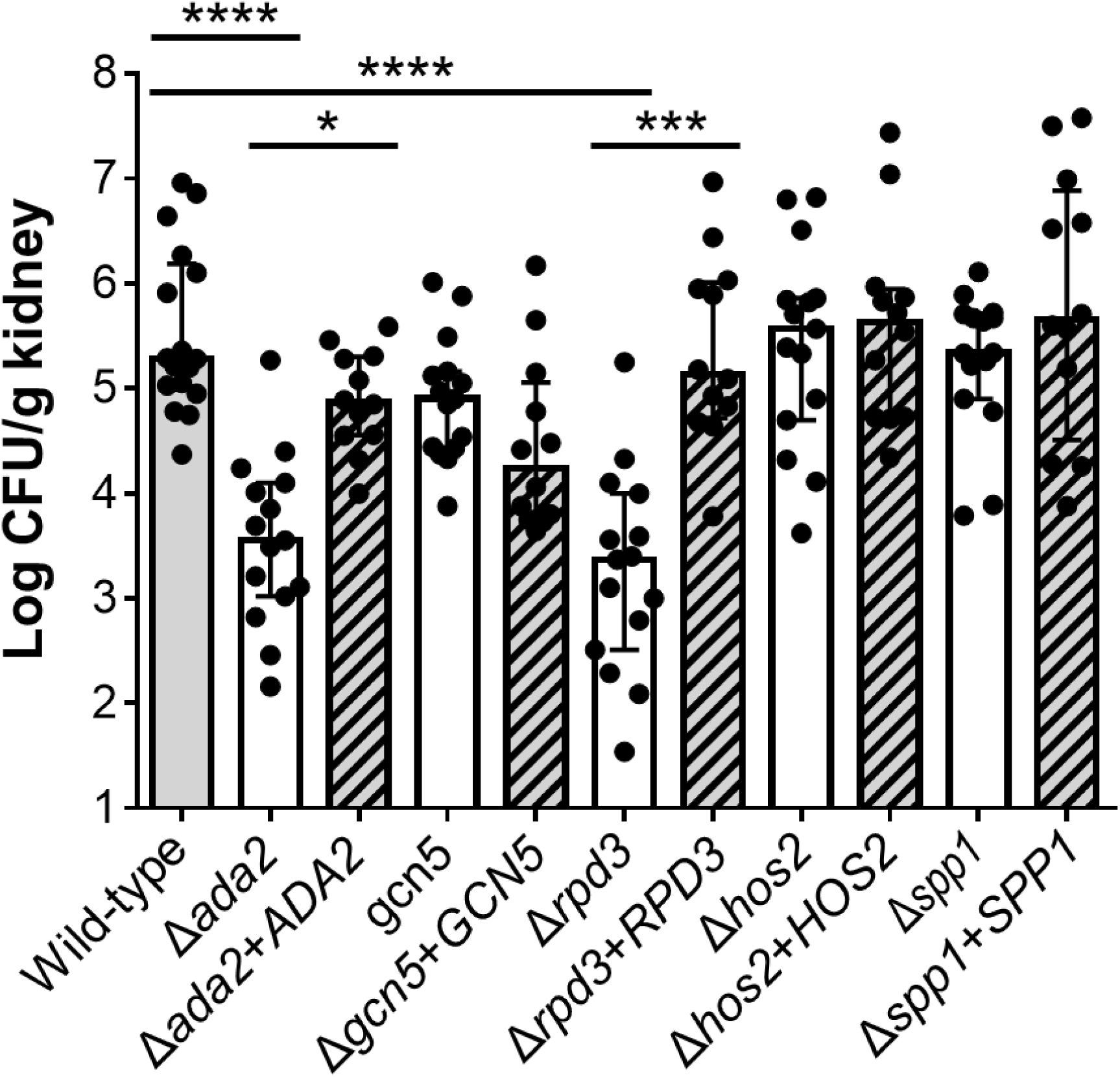
Virulence testing in mice. Neutropenic mice were inoculated intravenously with the indicated strains of *C. glabrata*. After 8 days, the mice were sacrificed and the kidney fungal burden was determined. Results are the median ± interquartile range of the combined results of two independent experiments for a total of 12-17 mice per strain. \**P* < 0.05, \*\*\**P* < 0.001, \*\*\*\**P* < 0.0001 by the Kruskal-Wallis test for multiple comparisons.

## Discussion

The results obtained from screening a library of *C. glabrata* deletion mutants revealed that histone modification plays a central role in governing the susceptibility of the organism to protamine and caspofungin. Of the eight transcriptional regulators that identified in this screen, six were predicted to be components of major regulatory complexes that modify histones. Furthermore, Ada2, an adapter protein that is a critical component of multiple histone acetyltransferase complexes, and RPD3, a histone deacetylase, are required for the maximal virulence of *C. glabrata* in neutropenic mice.

Ada2, Gcn5, and Spt8 were found to be necessary for *C. glabrata* to withstand both protamine and caspofungin. These proteins are members of the SAGA histone acetylase complex (11, 17). Ada2 functions as an adapter protein, Gcn5 is an acetyltransferase, and Spt8 is a regulatory protein that is required for the function of the SAGA complex (11). Ada2 and Gcn5 are also members of multiple other histone acetyltransferase complexes (18, 19). Although the phenotypes of the *ada2*Δ and *gcn5*Δ mutants were similar, the *ada2*Δ mutant had increased susceptibility to a broader range of stressors than the *gcn5*Δ mutant. For example, the *ada2*Δ mutant had increased susceptibility to calcofluor white and NaCl, whereas the *gcn5*Δ mutant did not. Also, while the *ada2*Δ mutant had a virulence defect in both *G. mellonella* and mice, the *gcn5*Δ mutant had only a modest virulence defect in *G. mellonella* and had wild-type virulence in mice. In *Candida albicans*, deletion of both *ADA2* and *GCN5* results in attenuated virulence (20, 21). Our results indicate that *C. glabrata* differs from *C. albicans* in that Ada2 plays a greater role than Gcn5 in stress resistance and virulence.

The finding that Ada2 is particularly important for the *C. glabrata* to tolerate a wide variety of stressors provides a compelling explanation as to why this mutant had reduced virulence in both *G. mellonella* and mice. Two other groups have also reported that *C. glabrata ada2*Δ mutants have increased susceptibility to a broad range of stressors (22, 23). In agreement with our results, Kounatidis et al. found that an *ada2*Δ mutant had attenuated virulence in a *Drosophila* model of gastrointestinal infection (22). By contrast, Yu et al. found that an *ada2*Δ mutant actually had increased virulence in mice, as manifested by enhanced lethality and increased organ fungal burden (23). The reason for why the virulence data of Yue et al. differed from ours is unclear, but may be due to differences in the background strain of *C. glabrata* and the immunosuppression regimen that were used.

We found that both the *rpd3*Δ and *hos2*Δ mutants has increased susceptibility to protamine and caspofungin, and that the *rpd3*Δ mutant had attenuated virulence in mice. In agreement with our results, Schwarzmuller et al. also determined that a *C. glabrata rpd3*Δ mutant had increased susceptibility to caspofungin (24). Rpd3 and Hos2 are members of the same class of histone deacetylases, and in *S. cerevisiae, rpd3*Δ and *hos2*Δ mutants have overlapping phenotypes (12). Other members of this class of histone deacetylases include Hda1, Hos1, and Hos3. We found that the *hda1*Δ mutant had increased resistance to protamine and wild-type susceptibility to caspofungin, and that the *hos1*Δ and *hos3*Δ mutants had wild-type susceptibility to both protamine and caspofungin. Thus, although these histone deacetylases are members of the same family, they have different functions in *C. glabrata*.

It was found that the *spp1*Δ mutant had increased susceptibility to protamine and caspofungin. Spp1 is predicted to be a member of the COMPASS complex. In *S. cerevisiae*, this complex functions as a histone methyltransferase, adding three methyl groups to lysine 4 of histone H3 by the Set1 methyltransferase. Spp1 stabilizes Set1 and retards its degradation (26). Our finding data suggest that the COMPASS complex is required for *C. glabrata* to withstand to protamine and caspofungin these stressors. The phenotype of a *C. glabrata spp1*Δ mutant has not been reported previously and a Δ*set1* mutant was not present in the library. However, it is known that a *C. albicans set1*Δ/Δ mutant is hyperfilamentous and has attenuated virulence in mice (27). Our experimental results predict that the Δ*set1* mutant would be likely susceptible to both protamine and caspofungin, and perhaps have attenuated virulence.

Testing the virulence of the various *C. glabrata* mutants in both *G. mellonella* and mice enabled us to determine the extent of concordance between these two models of disseminated candidiasis. The *ada2*Δ mutant had attenuated virulence in both models, and the *hos2*Δ and *spp1*Δ mutants had wild-type virulence in both models. By contrast, the *gcn5*Δ mutant had reduced virulence only in *G. mellonella*, and the *rpd3*Δ mutant had reduced virulence only in mice. These results suggest that although the *G. mellonella* model has modest predictive value, any *C. glabrata* mutant that has reduced virulence in this model should be tested in mice to verify the results.

Collectively, the data presented herein suggest that in *C. glabrata*, histone modification is a central mechanism by which the organism withstands stress due to host defense peptides and echinocandin antifungal drugs. Ada2 is also necessary for maximal virulence of this organism during disseminated infection. Although the transcriptional adapter-2α is the human counterpart of *C. glabrata* Ada2, these two proteins share only 29% amino acid identity. Thus, Ada2 holds promise as a potential antifungal drug target.

## Materials and Methods

### Deletion mutant library screening

To identify transcriptional regulators that govern resistance to protamine, a representative host defense peptide and caspofungin, a library of 215 *C. glabrata* transcriptional regulator deletion mutants constructed in the ATCC 2001 strain background was generously provided by Brendan Cormack at the Johns Hopkins School of Medicine (24). The susceptibility of the mutants to caspofungin and protamine was determined using an agar-dilution assay (28). The various mutants and the wild-type control strain were grown in yeast extract peptone dextrose (YPD) broth at 30°C in a shaking incubator overnight. The next day, each culture was adjusted to an OD_600_ of 2.7 in YPD broth. Four serial 10-fold dilutions were made in PBS, after which 3 µl of each dilution was spotted onto YPD agar containing either 325 ng/ml of caspofungin or 3 mg/ml of protamine. After incubation at 30°C for 48 h, the plates were imaged and the growth of each mutant was compared to that of the wild-type parent strain. Any mutant that differed from the wild-type strain in its susceptibility to caspofungin or protamine was independently tested at least twice to verify the results.

To assess whether selected mutants had increased susceptibility to additional stressors, the above agar dilution method was used to assess growth on calcofluor white (400 µg/ml), hydrogen peroxide (15 mM), SDS (0.2%), and NaCl (0.55 M).

### Complementation

To complement the Δ*ada2*, Δ*gcn5*, Δ*rpd3*, Δ*hos2*, and Δ*spp1* mutants, the protein coding regions plus 1000 bp of upstream and 500 bp of downstream flanking sequences of each gene was PCR amplified from *C. glabrata* genomic DNA. Each resulting PCR product was cut with BamHI and XhoI and ligated into plasmid pGRB2.1+His3 (29), which had been linearized with Bam HI and XhoI. The resulting plasmids were transformed into *C. glabrata* using the lithium acetate method.

### Radial diffusion assay

To assess the susceptibility of the wild-type, *ada2*Δ muant, and *ada2*Δ+*ADA2* complemented strains to human host defense peptides, we employed an ultrasensitive radial diffusion assay (30). The host defense peptides used were HNP-1 and hBD-2, and they were tested at pH 7.5.

### Virulence testing

The animal studies performed for this research were approved by the Lundquist Institute Animal Care and Use Committee. The virulence of the various *C. glabrata* mutants was assessed in the *G. mellonella* larval (31) and neutropenic mouse models of disseminated infection (32). *G. mellonella* larvae in the final instar stage were inoculated with 2.5 × 10^6^ or 5×10^6^ *C. albicans* cells in total volume of 5 μl using a Hamilton syringe. Twelve larvae were inoculated with each strain of *C. glabrata* and an additional 10 larvae were injected with 5μl of PBS as a negative control. The larvae were kept in a humidified incubator at 37°C and monitored twice daily for survival. Each strain was tested twice and the results of the two experiments were combined.

The virulence of the mutants was also tested in the neutropenic mouse model of disseminated candidiasis (32). To induce neutropenia, male Balb/c mice were administered 5-fluorouracil via intraperitoneal injection at 150 mg/kg of body weight at day −1 relative to infection. The next day, they were inoculated with 10^7^ *C. glabrata* cells via the lateral tail vein. After 8 days of infection, the mice were euthanized, after which their kidneys were harvested, weighed, homogenized, and quantitatively cultured. Each strain was tested twice in 6 to 8 mice per experiment and the results were combined.

### Statistical analysis

Using the GraphPad Prism software package, the survival of larvae infected with each mutant was compared with larvae infected with the wild-type strain and the complemented strain using the Gehan-Breslow-Wilcoxon test. The mouse organ fungal burden data were analyzed using the Kruskal Wallis test. P-values ≤ 0.05 were considered significant.

## Acknowledgements

We thank Brendan Cormack for generously supplying the transcription factor deletion library. This work was supported in part by NIH grant R01AI124566.

---

**Table S1. Results of screening the deletion library for susceptibility to protamine and caspofungin**.

